# Interaction of the Proline-Rich Domain of Fission Yeast WASp (Ws1p1) with Actin Filaments

**DOI:** 10.1101/2022.01.26.477833

**Authors:** Aaron D. Rosenbloom, Thomas D. Pollard

## Abstract

**Background:** The Wiskott-Aldrich Syndrome protein (WASp) family of proteins plays a crucial role in the activation of the Arp2/3 (actin-related protein 2/3) complex to promote the branching of actin filaments. The proline-rich domain (PRD) of WASp is known to contribute to branching nucleation but was overlooked, until experiments showed that the PRD of budding yeast Las17 can bind actin filaments (1).

**Methods:** We purified recombinant proline-rich domains from fission yeast *S. pombe* Wsp1 and budding yeast *S. cerevisiae* Las17 to test in biochemical assays of actin binding and polymerization.

**Results:** The PRD of the *S. pombe* Wsp1 binds actin filaments with micromolar affinity. The PRDs of both Wsp1 and Las17 slowed the rate of actin filament elongation by Mg-ATP-actin monomers by half and slowed the spontaneous polymerization of Mg-ATP-actin monomers modestly.

**Conclusion:** The affinity of PRDs of WASp-family proteins for actin filaments is high enough to contribute to the reported stimulation of actin filament branching by Arp2/3 complex.

## Introduction

Actin, one of the most prevalent proteins in eukaryotes, assembles into filaments that support the physical integrity and contribute to movements of cells (2). Spontaneous nucleation of actin filaments is unfavorable (3), but formins and Arp2/3 complex nucleate filaments at times and places appropriate for cellular functions (4). Formins nucleate linear filaments such as those found in the cytokinetic contractile ring of a dividing cell, while Arp2/3 complex produces branched filaments at the leading edge of motile cells to produce movements.

Arp2/3 complex consists of seven protein subunits (5) responsible for the nucleation of branched filaments (6). The two actin related proteins (Arps) share the same fold as actin but have different surface characteristics (7). These Arps form a nucleus that allows for branched filament formation (8). However, the native Arp2/3 complex is intrinsically inactive, owing to Arp2 and Arp3 being separated from each other, precluding the formation of the nucleus needed for actin polymerization (7). Proteins called nucleation-promoting factors (NPFs) and binding to the side of an actin filament activate Arp2/3 complex to form branches (9).

Nucleation promoting factors such as the Wiskott-Aldrich Syndrome protein (WASp) family have C-terminal motifs (10) including one or two verprolin-homology motifs (V) that bind actin monomers and central (C) and acidic (A) motifs that bind Arp2/3 complex (11). In addition to these VCA motifs, many NPFs have a proline-rich domain (PRD) before the VCA motif (12). The domains between the N-terminus and the PRD differ among the NPFs and contribute to regulation (12).

Studies of Arp2/3 complex activation by NPFs have focused on the VCA motif (13-15) but other evidence shows that the PRD plays a role in Arp2/3 complex activation (1, 9, 12, 16). First, the NPF activity of Scar, WASP, and Las17 are higher for constructs that include the PRD in addition to the VCA motif (9, 16, 17). Second, defects in endocytosis by budding yeast cells are more severe when both VCA and the PRD are removed from Las17, its WASp homolog (12). Third, the PRD of budding yeast Las17 was reported to stimulate the nucleation of actin filaments (1).

This evidence points to a role for the PRD in the activation of Arp2/3 complex, beyond its interaction with the SH3 domains of proteins that activate WASp and related proteins (18). However, much remained to be learned about the quantitative aspects of the molecular mechanisms. The work on Las17 opened a window into the role of the PRD, but other model organisms had not been tested. One such model organism is *S. pombe*.

*S. pombe* is a highly studied fission yeast (19) that has been used extensively to study endocytosis (20), a process driven by actin filament branching by Arp2/3 complex (21). Wsp1p is the WASp homologue of *S. pombe* (22). We measured affinity of Wsp1p-PRD for actin filaments and tested its effects on actin polymerization. We find that the PRD region binds actin filaments with micromolar affinity and slows the elongation of actin filaments by Mg-ATP-actin monomers.

## Methods and Materials

### Plasmid preparation

The DNA encoding amino acids 229 – 490 of *S. pombe* Wsp1p was amplified from genomic DNA and inserted into the pGEX-6P vector via In-Fusion cloning (Takara Bio USA). This vector added an N-terminal GST tag with a precision protease recognition site. After cleavage at this site, the vector leaves 5 amino acids on the N-terminus of the protein. The last of these five is amino acid 228, so the final protein includes amino acids 228-490. Six histidine residues were added to the C-terminus of the sequence via the Q5 Site-Directed Mutagenesis Kit (New England Biolabs). The resulting vector was transformed into OverExpress C41(DE3) chemically competent *E. coli* cells (Lucigen). To allow for fluorescent labeling of Wsp1p-PRD, a cysteine was added to the N-terminus of the coding sequence directly after the precision protease recognition site or after the six-histidine tag on the C-terminus using the Q5 Site-Directed Mutagenesis Kit (New England Biolabs).

A plasmid encoding amino acids 301-536 of *S. cerevisiae* Las17 in a pGEX-6P vector was a generous gift from the laboratory of Kathryn Ayscough at the University of Sheffield. This plasmid attaches a GST-tag to the N-terminus of Las17-PRD. When cleaved by lab made Precision Protease, five amino acids are left at the N-terminus of the PRD.

### Protein Expression and Purification

Transformed *E. coli* were grown in flasks with 1 L of LB-ampicillin at 37 °C with shaking at 200 RPM. When the optical density at 595 nm reached 0.4 – 0.5, we added 100 μL of 1 M isopropyl ß-D-1-thiogalactopyranoside (IPTG) and incubated overnight at 18 °C with shaking at 250 RPM. The cells were harvested by centrifugation at 5,000 g for 15 min. Bacterial pellets from 4 L of cell culture were resuspended in 165 mL of lysis buffer at pH 7.4 (480 mM NaCl, 3 mM KCl, 10 mM phosphate, 10 mM imidazole, 1 mM PMSF, and an inhibitor cocktail that gave final concentrations of 1 μg/mL of bestatin, aprotinin, leupeptin, and pepstatin). The resuspended cells were frozen and stored at -20 °C until use.

Resuspended pellets of cells from 4 L of culture were thawed in a room temperature water bath and sonicated on ice 8 times for 1 min with one-minute rest intervals using a 3/8 inch probe at 50% duty cycle and an output control of 6 on a Branson Sonifier 450. The lysate was clarified by centrifugation in a Type 45 Ti rotor (Beckman Coulter) at 35,000 RPM for 35 min at 4 °C. The supernatant stirred with 10 mL of TALON metal affinity resin (Takara Bio US) equilibrated in Buffer A (480 mM NaCl, 3 mM KCl, 10 mM phosphate, and 10 mM imidazole, pH 7.4) for 1 h at 4 °C. The resin was pelleted and washed three times with 40-45 mL of cold wash buffer. Bound protein was eluted from the washed resin four times for at least 10 min with 10 mL of cold elution buffer at a pH of 7.4 (Buffer A with an additional 240 mM imidazole) followed by pelleting after each elution. The combined eluants were incubated overnight at 4 °C with 5 mL of Glutathione-Sepharose 4B resin (GE Healthcare Bio-sciences AB). The beads were placed in a column and washed 3 times with 50 mL of Buffer B (480 mM NaCl, 3 mM KCl, and 10 mM phosphate, pH 7.4) and three times with 50 mL of KMEID buffer (50 mM KCl, 1 mM MgCl_2_, 1 mM EGTA, 10 mM imidazole pH 7.0, 2 mM DTT). The beads were suspended in 5 mL of KMEID and the protein was cleaved overnight at 4 °C using ∼0.2 mg of lab made GST tagged Precision Protease. The cleaved protein was collected from the column and concentrated approximately 10-fold using an Amicon Ultra Centrifugal Filter Unit (Millipore) with a 3,000 Da cutoff. The concentrated protein was dialyzed two times against 500-1,000 mL KMEI buffer.

For labeling with Alexa488, Wsp1p-PRD and Wsp1p-PRD were purified up to the first set of washes of the Glutathione-Sepharose 4B resin. The beads were then washed with 5 mL of KMIT (50 mM KCl, 1 mM MgCl_2_, 10 mM imidazole pH 7.0, 2 mM tris(2-carboxyethyl)phosphine, TCEP) and the protein was cleaved with lab made GST tagged Precision Protease as above. The cleaved protein was labeled by introducing 500 nmole Alexa Fluor 488 maleimide (Thermo Fisher Scientific) to the cleaved solution and incubating at 4 °C overnight. The conjugated protein was incubated with 200 μL of equilibrated TALON resin for one hour. The beads were pelleted and washed 3 times with 1 mL of KMIT buffer. The protein was eluted with 5 sets of 200 μL KMI+ (50 mM KCl, 1 mM MgCl_2_, 250 mM imidazole pH 7.0). The protein was dialyzed against two changes of 500 mL of KMEI and concentrated using an Amicon Ultra Centrifugal Filter Unit (MilliporeSigma) with a 3,000 Da cutoff. The concentration of the dye conjugated to the protein was determined by absorption at 495 nm with an extinction coefficient of 71,000 M^-1^ cm^-1^.

Las17-PRD was expressed in OverExpress C41(DE3) *E. coli* cells (Lucigen) and purified as described by Urbanek *et al*. (1) with minor modifications. One-liter cultures of TB-ampicillin were incubated at 37 °C with shaking at 200 RPM until an optical density at 595 nm of approximately 0.4 – 0.5. Then each flask was induced with 1 mL of 1 M IPTG overnight at 37 °C with shaking at 250 RPM. Cell pellets were resuspended in 40 mL of modified lysis buffer (10 mM phosphate buffer pH 7.4, 480 mM NaCl, 3 mM KCl, 1 mM PMSF, and an inhibitor cocktail that gave final concentrations of 1 μg/mL of bestatin, aprotinin, leupeptin, and pepstatin) per 4 L of culture. The resuspended cells were frozen and stored at -20 °C until use.

Pellets from 4 L of culture were thawed at room temperature, sonicated on ice and clarified as above. The supernatant was added to 5 mL of equilibrated Glutathione Sepharose 4B resin (GE Healthcare Bio-sciences AB). The beads were washed in a column 3 times with 50 mL of Buffer C (10 mM phosphate buffer pH 7.4, 480 mM NaCl, 3 mM KCl, and 0.1% Tween-20), three times with 50 mL of Buffer D (10 mM phosphate buffer pH 7.4, 140 mM NaCl, and 3 mM KCl), two times with 50 mL of Buffer E (50 mM Tris-HCl, 400 mM NaCl, 1 mM EDTA, 1 mM DTT, pH 7) and two times with 50 mL of Buffer F (50 mM Tris-HCl, 150 mM NaCl, 1 mM EDTA, 1 mM DTT, pH 7). The beads were placed in 5 mL of Buffer F and the protein was cleaved overnight at 4 °C using ∼0.2 mg of lab made GST tagged Precision Protease. The cleaved protein was removed from the column, concentrated, and dialyzed two times against 500 – 1,000 mL of KMEI buffer as above.

### Actin purification and labeling

Actin was isolated from an acetone powder of chicken breast muscle and purified by one cycle of polymerization, pelleting, and depolymerization before gel filtration chromatography on a Sephacryl-S300 column in G-buffer (23). For labeling, actin filaments at a concentration of 1 mg/mL in 0.1 M KCl, 2 mM MgSO_4,_ 25 mM Tris pH 7.5, 3 mM NaN_3_, and 0.3 mM ATP were reacted with a 10-fold molar excess of *N*-(1-pyrene)iodoacetamide (Thermo Fisher Scientific) dissolved in dimethylformamide (DMF) at 4 °C overnight. The actin filaments were pelleted, homogenized in G-buffer, depolymerized by dialysis against G-buffer for two days at 4°C, and clarified at 75,000 RPM in a TLA 120.2 rotor (Beckman Coulter) at 4 °C for 2 h. The upper two thirds of the supernatant were gel filtered on a Superdex-200 column in G-buffer.

### Mass spectrometry

A sample of the purified protein was run on a Mini-PROTEAN TGX Stain-free gel (BIORAD) with 4-20% acrylamide gradient and stained with Coomassie brilliant blue. The stained bands were cut out of the gel. The Mass Spectrometry and Proteomics Resource of the W. M. Keck Foundation Biotechnology Resource Laboratory of Yale University digested the bands with trypsin and analyzed the peptides by MS/MS mass spectrometry.

### Concentration determination

The concentrations of samples of Wsp1p-PRD and Las17-PRD were determined by SDS-PAGE, staining with Coomassie brilliant blue and densitometry with a range of amounts of chicken skeletal muscle actin as the standard. Samples were run on a 15-well Mini-PROTEAN TGX Stain-free gel (BIORAD) with 4-20% polyacrylamide gradient. Gels were stained with Coomassie blue, destained 3 times for 10 min with 25% methanol and 10% acetic acid, and soaked in water overnight. Gels were imaged with a ChemiDoc Imaging System (BIORAD). The densities of the bands were determined with ImageJ software. The mass of the PRD sample was determined by comparison with the linear standard curve. A molar concentration of the protein was calculated using molecular weights of 26.7 kD for Wsp1p-PRD and 24.6 kD for Las17-PRD.

### Actin filament binding assay

A high concentration of actin monomers was polymerized in KMEI buffer and used to prepare 100 μL samples with 0, 2.5, 5, 10, 25, 50, 100, and 180 μM actin and 1 μM of Alexa488-Wsp1p-PRD or Wsp1p-PRD-Alexa488. Samples were centrifuged in a TLA 100 rotor (Beckman Coulter) at 75,000 RPM for 30 min. The fluorescence of 75 μL of supernatant was measured with a Gemini EM microplate reader (Molecular Devices) with excitation at 495 nm and emission at 519 nm with a cutoff at 515 nm with 6 flashes per read. The plate was turned 180 degrees and re-measured to be certain there was no instrumental bias in the measurements.

The fluorescence of the supernatants of the 50 μM, 100 μM, and 180 μM actin samples were averaged and used to correct for background fluorescence. The corrected fluorescence measurements of the 0 μM actin samples were averaged to determine the value of unbound protein. Subtracting the ratio of each corrected fluorescence value to the average fluorescence value of 100% unbound protein from 1 gave the fraction of bound protein. The fraction of bound protein was plotted against the concentration of actin. The dissociation equilibrium constant (K_d_) was determined by fitting 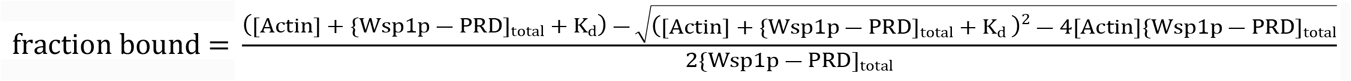 to the data.

### Fluorescence anisotropy experiments

Concentrations of 20 nM Alexa488-Wsp1p-PRD or 100 nM Wsp1p-PRD-Alexa488 were incubated in KMEI buffer at room temperature for 30 min with 0, 2.5, 5 or 10 μM filamentous actin. Anisotropy was measured with an Alpha-scan spectrofluorometer (Photon Technology International) with excitation at 495 nm and emission of 519 nm. The excitation bandpass was set to 4 nm and the emission bandpass was at 20 nm. Each sample was measured at four different settings: first (HV), excitation polarizer at 90° and the emission at 90°; second (HH), excitation at 90° and emission at 0°; third (VV), excitation at 0° and emission at 0°; and fourth (VH), excitation polarizer at 0° and emission at 90°. Emission measurements were taken every second for 10 s. For Alexa488-Wsp1p-PRD 5 replicates were collected for 2.5, 5, and 10 μM polymerized actin, and a total of 6 replicates for 0 μM filamentous actin. For Wsp1p-PRD-Alexa488, only one sample was measured for each concentration.

To calculate the anisotropy of each sample, the 10 measurements of HH, HV, VV, and VH were averaged for each sample. The G-factor for each sample was calculated according to 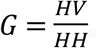. All the calculated G-factors were averaged together to determine the final G-factor. The anisotropy was calculated according to the formula 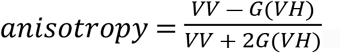.

### Assay for the polymerization of bulk samples of actin

Pyrene-labeled (10%) Ca-ATP-actin monomers were exchanged for Mg^2+^ by adding 10x MEA buffer resulting in 50 μM MgCl_2_, 200 μM EGTA pH 8.0, 0.00005% Antifoam 204 (Sigma) and incubating at room temperature for 2-5 min. Polymerization was initiated by mixing two volumes of PRD in 1.5x of KMEIA buffer (1x: 50 mM KCl, 1 mM MgCl_2_, 1 mM EGTA, 10 mM imidazole pH 7.0, 0.00005% Antifoam 204) or KMEIA alone with the Mg-ATP-actin monomers. The time course of the fluorescence change was measured with a SpectraMax Gemini EM Microplate Reader (Molecular Devices) with excitation at 365 nm and emission at 407 nm in a Costar 3694 EIA/RIA, 96 well, half area, flat bottom, non-treated, black polystyrene plates using one flash per read every 2 min for 1 μM and 2.5 μM actin, every 10 s for 5 μM and 7.5 μM actin, and every 5 s for 10 μM actin. Since the 1 μM and 2.5 μM samples were run overnight, an optical adhesive was placed on top to prevent evaporation.

The data was processed by adding the deadtime and normalized similar to Doolittle *et al*. (24) for the initial minimum (I_min_) and final maximum (I_max_) values. Fluorescence intensities were converted to the polymerized actin concentration by first subtracting the I_min_ from each intensity and taking into account the critical concentration of 0.1 μM.

### Measurement of elongation rates

Samples of 150 μL in wells of Costar 3694 EIA/RIA polystyrene plates consisted of 2 μM Mg-ATP-actin monomers, 0.5 μM pre-polymerized actin filament seeds, and either no PRD, 7.5 μM Wsp1p-PRD or 7.5 μM Las17-PRD in 50 mM KCl, 1 mM MgCl_2_, 1 mM EGTA, 10 mM imidazole pH 7.0, 0.00005% Antifoam 204. Fluorescence was measured every 5 s with a SpectraMax Gemini EM Microplate Reader (Molecular Devices) with excitation at 365 nm and emission at 407 nm. The dead time between adding actin monomers and acquisition was noted.

The data was processed similar to the spontaneous polymerization data. To correct for the photobleaching, the intensities after T_max_ for the no PRD samples were fit to an exponential. The exponentials for all the samples without PRD were averaged together (n = 6) and used to correct for photobleaching. The samples without PRD were used to determine I_min_ and I_max_ for all samples. We calculated the number concentration of filaments for the reactions without PRD from the rate of elongation and the rate constant for elongation (25). To calculate the barbed-end elongation rates for each of the reactions, we divided the average rate of polymerization of each set of samples in μM s^-1^ by the concentration of filament ends. We measured a total of six replicates without PRD, four replicates for Wsp1p-PRD, and three replicates for Las17-PRD.

## Results

Attempts to purify Wsp1p-PRD with just a GST tag at the N-terminus produced many protein fragments, likely due to ribosome stalling on polyproline sequences (26) and producing truncated proteins. To eliminate these fragments, we appended a 6x-histidine tag to the C-terminus of GST-Wsp1p-PRD. After two affinity purification steps, Wsp1p-PRD ran as the largest band on SDS-PAGE with only slight contamination (Fig. 1). Mass spectrometry of a trypsin digest of the major band confirmed it is Wsp1p-PRD. The Wsp1p-PRD band ran slower than expected from its molecular weight of 26.7 kD (Fig. 1), similar to other proline-rich proteins (27). We also purified Las17-PRD by small modifications of the method of Urbanek *et al*. (1) (Fig. 2). Mass spectrometry of a trypsin digest identified the dominant band as Las17-PRD, the higher molecular weight contaminant as the *E. coli* chaperonin dnaK and the minor lower molecular weight bands as fragments of Las17-PRD, likely produced by ribosomal stalling.

**Fig. 1.**
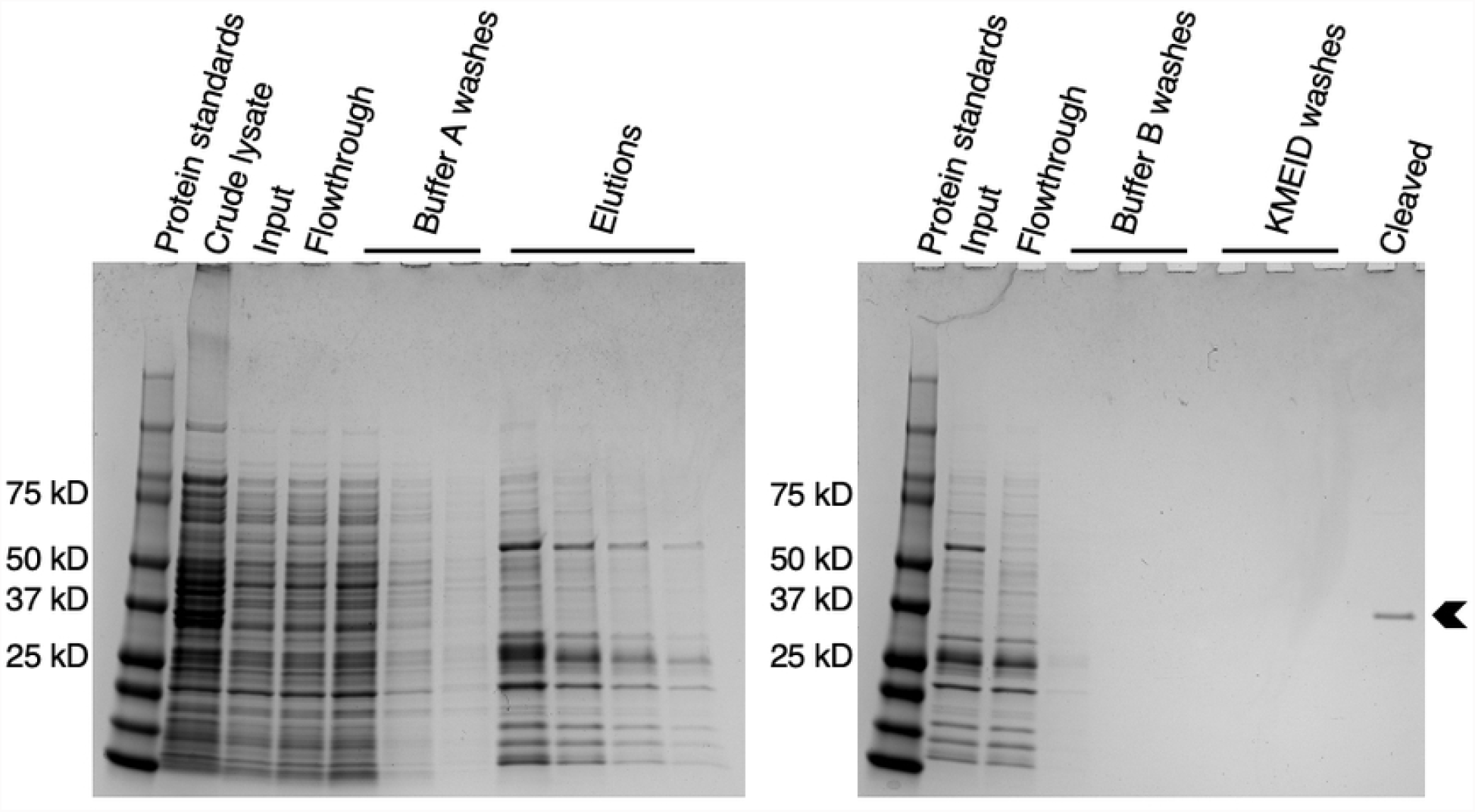
SDS-PAGE and Coomassie brilliant blue staining of samples from each step of the purification of Wsp1p-PRD. Left gel: lane 1, protein standards; lane 2, crude lysate; lane 3 clarified sample of crude lysate applied to TALON resin; lane 4, unbound flowthrough from TALON resin; lanes 5-7, successive washes of TALON resin with Buffer A (480 mM NaCl, 3 mM KCl, 10 mM phosphate, 10 mM imidazole pH 7.4); lanes 8-11, elution of bound proteins from TALON resin with elution buffer (Buffer A with an additional 240 mM imidazole pH 7.4). Right gel: lane 1, protein standards; lane 2, combined fractions eluted from TALON resin and applied to glutathione Sepharose 4B beads; lane 3, unbound flowthrough from glutathione Sepharose 4B beads; lanes 4-6, successive washes of the glutathione Sepharose 4B beads with Buffer B (480 mM NaCl, 3 mM KCl, and 10 mM phosphate at a pH 7.4); lanes 7-9, washes of glutathione Sepharose 4B beads with KMEID (50 mM KCl, 1 mM MgCl_2_, 1 mM EGTA, 10 mM imidazole pH 7.0, 2 mM DTT); lane 10, pure Wsp1p-PRD cleaved from glutathione Sepharose 4B beads. The arrowhead marks Wsp1p-PRD.

**Fig. 2.**
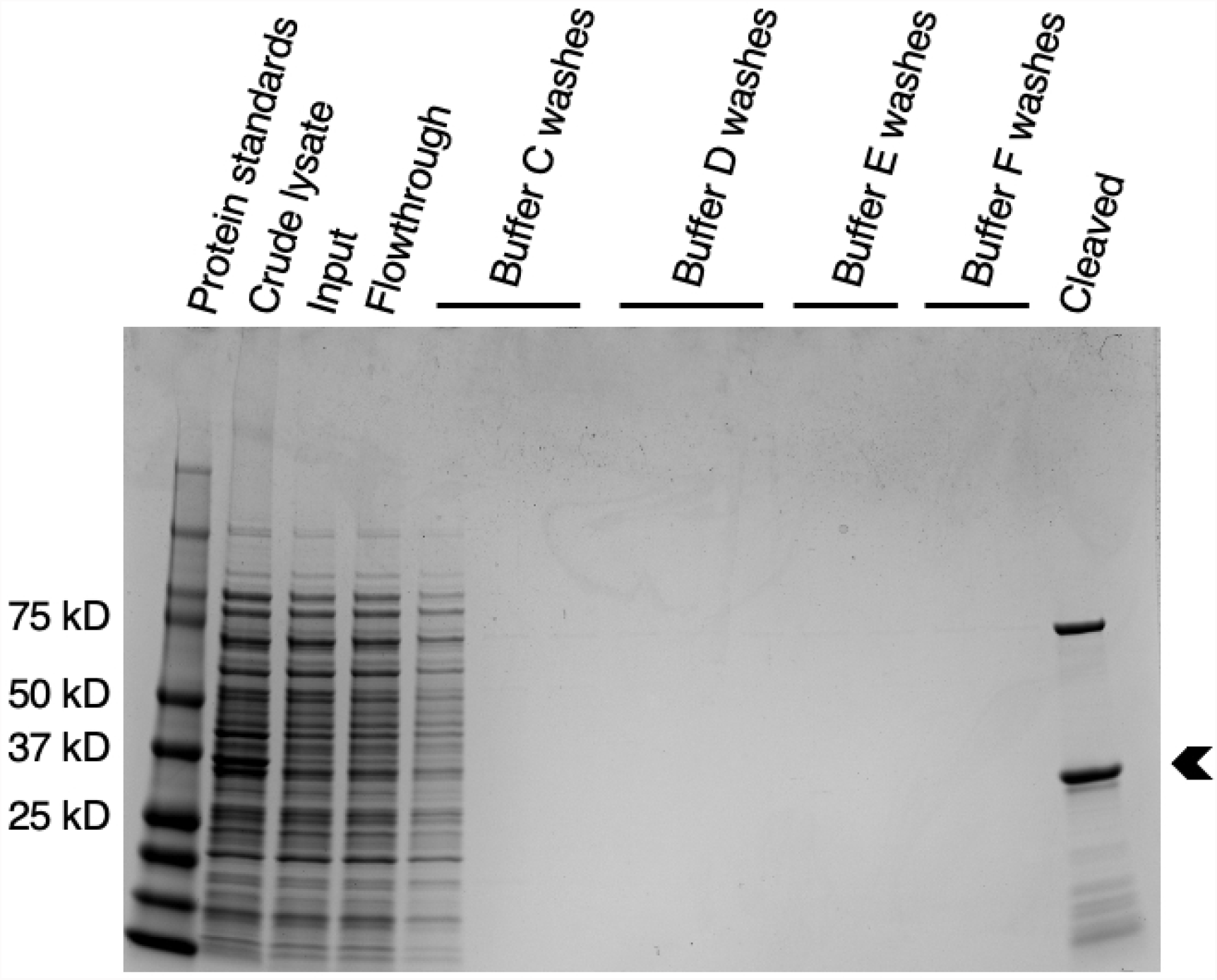
Coomassie brilliant blue stained SDS-PAGE gel of purification steps of *S. cerevisiae* Las17-PRD. Lane 1, protein standards; lane 2, crude lysate; lane 3, clarified lysate applied to glutathione Sepharose 4B beads; lane 4, unbound flowthrough from glutathione Sepharose 4B beads; lanes 5-7, washes of glutathione Sepharose 4B beads with Buffer C (10 mM phosphate buffer pH 7.4, 480 mM NaCl, 3 mM KCl, and 0.1% Tween-20); lanes 8-10, washes of glutathione Sepharose 4B beads with Buffer D (10 mM phosphate buffer pH 7.4, 140 mM NaCl, and 3 mM KCl); lanes 11-12, washes of glutathione Sepharose 4B beads with Buffer E (50 mM Tris-HCl, 400 mM NaCl, 1 mM EDTA, 1 mM DTT, and brought to a pH of 7 at room temperature); lanes 13-14, washes of glutathione Sepharose 4B beads with Buffer F (50 mM Tris-HCl, 150 mM NaCl, 1 mM EDTA, 1 mM DTT brought to pH 7 at room temperature); lane 15, protein cleaved from glutathione Sepharose 4B beads. Mass spectrometry of tryptic digests identified the main band (arrowhead) as Las17-PRD, the upper band as *E. coli* dnaK, and the minor lower molecular weight bands as fragments of Las17-PRD, likely due to ribosome stalling.

PRDs have no aromatic residues to measure absorbance at 280 nm; therefore, we estimated their concentrations by densitometry of Coomassie brilliant blue stained Wsp1p-PRD bands run on SDS-PAGE compared with a standard curve of known amounts of actin. This measurement is not the ideal, as proteins may stain differently in Coomassie blue (28, 29).

We determined the binding affinity of fluorescently labeled Wsp1p-PRD for actin filaments by measuring the fraction of 1 μM N-terminally (Alexa488-Wsp1p-PRD) or C-terminally (Wsp1p-PRD-Alexa488) labeled Wsp1p-PRD that pelleted with a range of actin filament concentrations from 0 – 180 μM (Fig. 3A and B). We calculated the fraction bound by measuring the fluorescence of unbound Wsp1p-PRD in the supernatants. Fitting a binding isotherm to the data gave a K_d_ of 6.2 ± 0.6 μM for Alexa488-Wsp1p-PRD (Fig. 3A). We confirmed that the N-terminal fluorescent tag does not affect binding by measuring the fraction for C-terminally (Wsp1p-PRD-Alexa488) labeled Wsp1p-PRD that pelleted with a range of actin filament concentrations (Fig. 3B). The K_d_ of 5 ± 1 μM for Wsp1p-PRD-Alexa488 was similar to Alexa488-Wsp1pPRD.

**Fig. 3.**
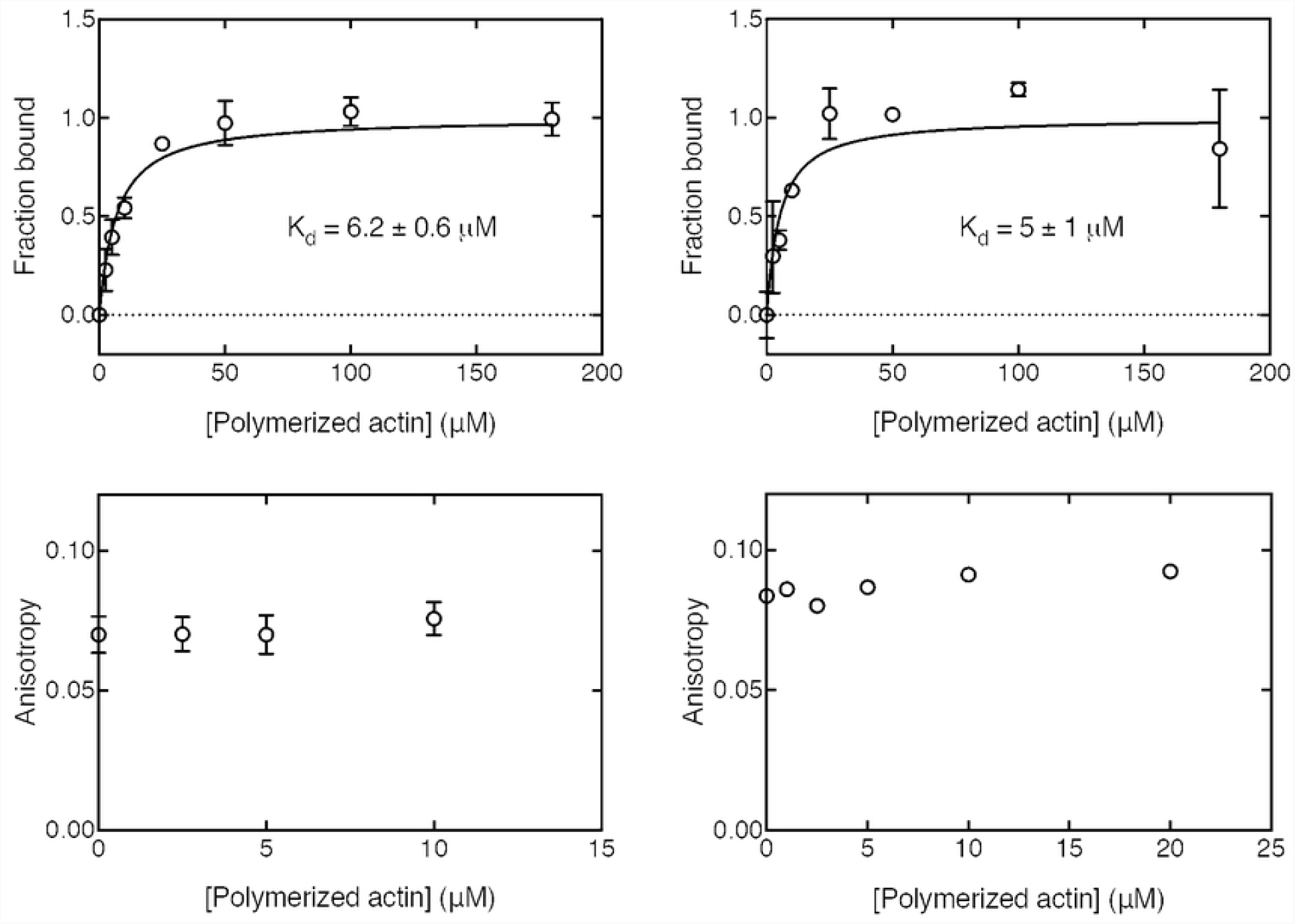
Affinity of labeled Wsp1p-PRD for actin filaments determined by cosedimentation and fluorescence anisotropy. (A-B) A range of concentrations of actin filaments were incubated with 1 μM of Alexa-488 labeled Wsp1p-PRD in 50 mM KCl, 1 mM MgCl_2_, 1 mM EGTA, 10 mM imidazole at pH 7.0 for 30 min at 25°C followed by ultracentrifugation at 240,000 g for 30 min. The fraction bound was calculated from the fluorescence of the supernatant. (A) Fraction of Alexa488-Wsp1p-PRD bound vs. actin filament concentration. Points are the mean with standard deviation as the error (n = 5-6). The solid line is the fit to the data of a binding isotherm with a K_d_ of 6.1 ± 0.6 μM. (B) Fraction of Wsp1p-PRD-Alexa488 bound vs. actin filament concentration. Points are the mean with the range as the error (n = 2-3). The solid line is the fit to the data of a binding isotherm with a K_d_ of 3 ± 1 μM. (C-D) A range of concentrations of actin filaments were incubated with 20 nM of Alexa488-Wsp1p-PRD or 100 nM of Wsp1p-PRD-Alexa 488 in 50 mM KCl, 1 mM MgCl_2_, 1 mM EGTA, and 10 mM imidazole at pH 7.0 at 25°C. Each sample was excited with polarized light, and the anisotropy was calculated based on the polarized components of the emission. (C) Fluorescence anisotropy of Alexa488-Wsp1p-PRD vs. concentration of actin filaments. Points are the mean ± standard deviation (n = 5-6). (D) Fluorescence anisotropy of Wsp1p-PRD-Alexa488 vs. concentration of actin filaments. Points are values from one experiment.

We used similar samples to measure the effects of actin filaments on the fluorescence anisotropy of the N- and C-terminally tagged Wsp1p-PRD. Surprisingly, actin filaments did not change the anisotropy in spite of binding the tagged Wsp1p-PRD (Fig. 3C and D). Therefore, binding to actin filaments did not change the rotational diffusion of Alexa-488 at either end of Wsp1p-PRD.

The presence of 5 μM Wsp1p-PRD or Las17-PRD slowed the time course of spontaneous polymerization of 5 μM Mg-ATP actin monomers modestly (Fig. 4A). This is the opposite to the previous reports for 0.08-0.14 μM Las17-PRD with 5 μM rabbit muscle Ca-ATP-actin in polymerization buffer with MgCl_2_ and EGTA at pH 8.0 (1, 30).

**Fig. 4.**
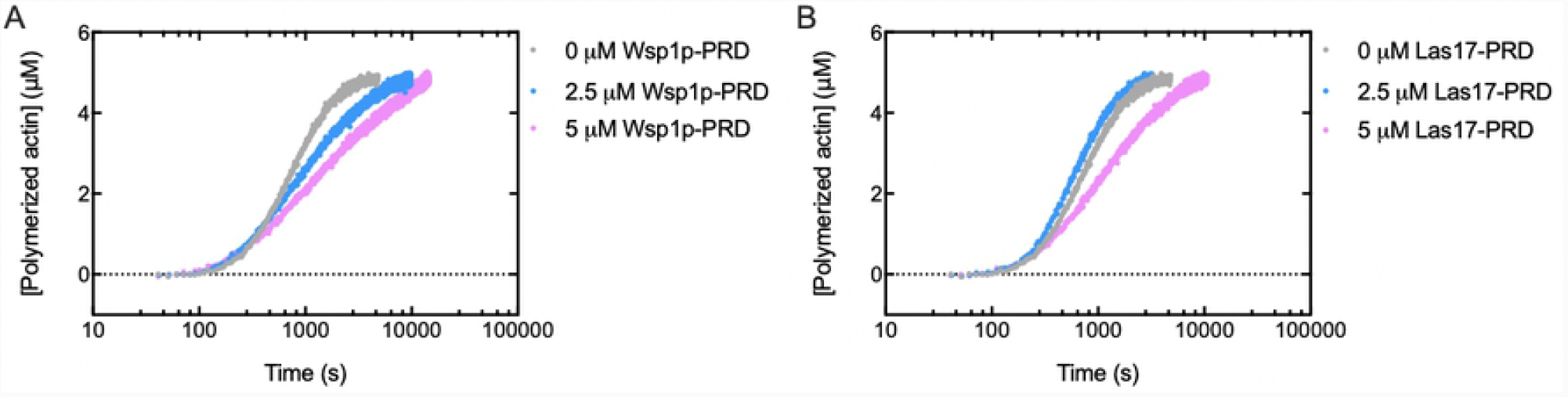
Effects of PRDs on the time course of the spontaneous polymerization of 5 μM of Mg-ATP-actin monomers (10% pyrenyl-actin). Conditions: 50 mM KCl, 1 mM MgCl_2_, 1 mM EGTA, 10 mM imidazole pH 7.0, 0.6 mM Tris, 0.15 mM DTT, 60 μM ATP, 30 μM CaCl_2_ and 0.00005% Antifoam 204 in the presence of two concentrations of (A) Wsp1p-PRD or (B) Las17-PRD.

To understand how the PRD slows spontaneous polymerization, we used the fluorescence of pyrenyl-actin to measure the elongation rates of actin filament seeds in the presence of PRD. The polymerization samples contained 2 μM Mg-ATP-actin (10% labeled with pyrene), 0.5 μM of polymerized actin to provide seeds, and either no PRD, 5 μM Wsp1p-PRD, or 5 μM Las17-PRD (Fig. 5A). The fluorescence increased linearly between 50 and 120 s (Fig. 5B) characteristic of pure elongation without a contribution from spontaneous nucleation (31). Polymerization of 2.5 μM Mg-ATP-pyrenyl-actin monomers without filament seeds started with a clear lag (Fig. 5A). Thus, under the conditions used, little spontaneous nucleation occurred during the first 120 s of the reaction, so all of the polymerization during this time came from elongation of previously formed filament seeds.

**Fig. 5.**
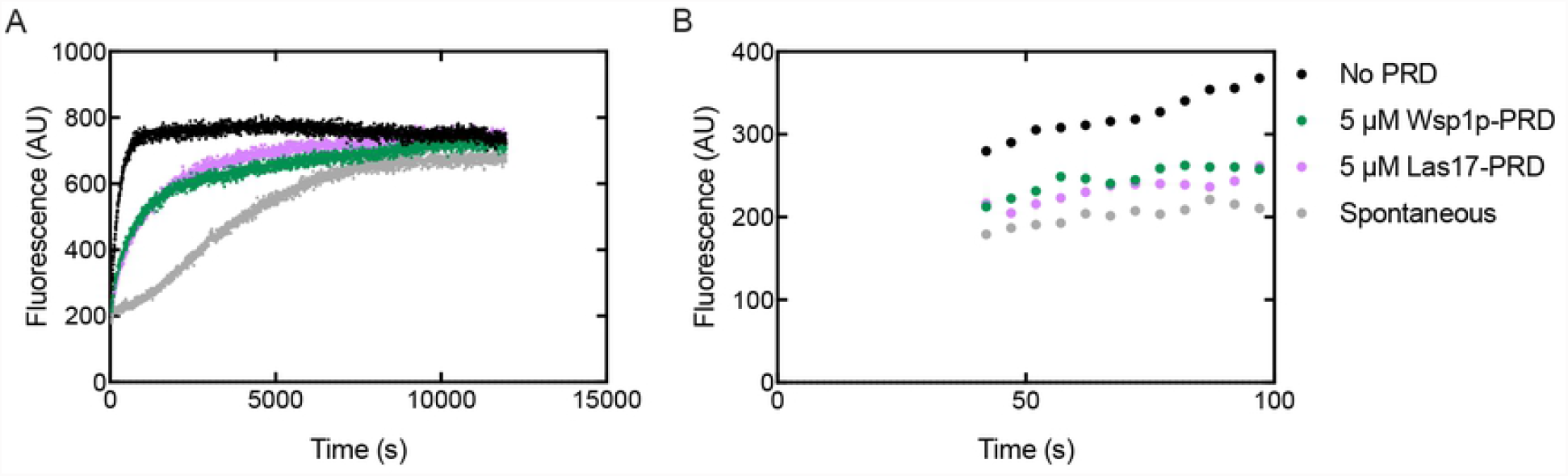
Effect of PRDs on the time course of elongation of actin filaments from preformed seeds (0.5 μm polymerized actin, 0.3 nM ends). Conditions: 2 μM 10% Mg-ATP-pyrenyl-actin monomers with either no PRD (black), 5 μM Wsp1p-PRD (green), or 5 μM of Las17-PRD (pink). For comparison, 2.5 μM of 10% Mg-ATP-pyrenyl-actin with the equivalent amount of labeled actin were polymerized without preformed filaments or PRD (grey). (A) Examples of time courses corrected for photobleaching. (B) Initial parts of the polymerization curves from A after the initial dead time.

Since the rate constants for actin filament elongations are known (25) we used the samples without PRD to calculate that the concentration of filament ends added to each reaction was 300 ± 100 pM (average concentration ± standard deviation; n = 6). The rate of barbed end elongation of samples of 2 μM actin monomers was 20 ± 10 subunits s^-1^ without PRD; n = 6 for rate), 10 ± 4 subunits s^-1^ with 5 μM of Wsp1p-PRD (same as above; n = 4 for rate), and 9 ± 3 nM s^-1^ with 5 μM of Las17-PRD (same as above; n = 3 for rate). Therefore, both Wsp1p-PRD and Las17-PRD reduced the elongation rate by approximately half.

## Discussion

### Binding affinity

Previous work (1) presented evidence that the PRD of Wsp1p family proteins can bind actin filaments but did not measure the affinity of the interaction with standard methods. The methods used included two-hybrid assays, pelleting with single concentrations of PRD, and actin and polymerization time courses. Urbanek *et al. (1)* estimated the affinity for actin filaments to be 0.22 μM from a double reciprocal plot of 1/actin polymerization rate vs. 1/[Las17 PRD] (their Fig. S2), but <20% of 5 μM PRD pelleted with 5 μM actin filaments corresponding to a K_d_ of about 20 μM (their Fig. 1D). Microscale thermophoresis was also used to measure a K_d_ of 24 nM for the 300–422 fragment of Las17 binding to actin monomers (1, 30).

Our quantitative binding assays confirmed that the Wsp1-PRD binds muscle actin filaments with micromolar affinity using Wsp1-PRD labeled on either end with Alexa-488. Surprisingly, even when most of the Wsp1-PRD was bound to the filaments, the Alexa-488 on both ends of the PRD remained mobile. This is consistent with binding being mediated by the middle of the PRD as shown for Las17-PRD (1, 30) and the known flexibility of peptides with disordered links between proline-rich sequences (32).

### Effects on polymerization

Urbanek *et al*. (1) reported that submicromolar concentrations of Las17 PRD stimulate spontaneous polymerization of 5 μM Ca-ATP-actin monomers in buffer with MgCl_2_ (conditions where the exchange of bound Ca^2+^ for Mg^2+^ is rate limiting), consistent with promoting nucleation but also with an effect on divalent cation exchange. They also reported that submicromolar concentrations of Las17-PRD increased the steady-state fluorescence of pyrenyl-actin filaments, which they interpreted as a decrease in the critical concentration. However, a substantial change in the polymer concentration under their conditions without PRD would have been unlikely, since the polymer concentration would have been 4.9 μM and the monomer concentrations 0.1 μM, an already low concentration.

We found that 5 μM concentrations of either Wsp1p-PRD or Las17-PRD slow the time course of spontaneous polymerization of 5 μM Mg-ATP-actin monomers, while 2.5 μM Las17-PRD had less effect on spontaneous polymerization. Both Las17-PRD and Wsp1p-PRD slowed the elongation of actin filament barbed ends by about half. This is a plausible explanation for the inhibition of spontaneous polymerization; however, more work is necessary to fully understand the mechanism of inhibition. This new finding raises the possibility that the effect of Las17-PRD on the polymerization of Ca-ATP-actin in buffer containing MgCl_2_ (1) may have been due to increasing the rate limiting exchange of Mg^2+^ for Ca^2+^ rather than effect on the polymerization reaction.

The micromolar affinity of Wsp1p-PRD for actin filaments has interesting implications for its function. As the total concentration of actin in the fission yeast cell is >50 μM and much higher than this average in endocytic actin patches (33), it stands to reason that a substantial fraction of Wsp1 is bound to actin filaments inside cells, especially at sites where branched actin filaments form. Given the mobility of both ends of the PRD bound to an actin filament revealed by fluorescence anisotropy (Fig. 3C and D), the VCA region of a Wsp1 molecule bound by its PRD to an actin filament should be capable of binding and activating Arp2/3 complex to nucleate an actin filament branch on a mother filament. On the other hand, the 4 μM concentration of Wsp1 in the cytoplasm of fission yeast (34) will likely have a only a small effect on the growth of the much higher concentration of actin filaments.

## Acknowledgements

Research reported in this publication was supported by National Institute of General Medical Sciences of the National Institutes of Health under award number R01GM026338. The content is solely the responsibility of the authors and does not necessarily represent the official views of the National Institutes of Health. The authors thank the laboratory of Kathryn Ayscough at the University of Sheffield for providing the plasmid for Las17-PRD and advice about its purification.

## Competing interests

The authors declare that no competing interests exist.

## Notes

### Competing Interest Statement

The authors have declared no competing interest.

